# Channel Embedding for Informative Protein Identification from Highly Multiplexed Images

**DOI:** 10.1101/2020.03.24.004085

**Authors:** Salma Abdel Magid, Won-Dong Jang, Denis Schapiro, Donglai Wei, James Tompkin, Peter K. Sorger, Hanspeter Pfister

## Abstract

Interest is growing rapidly in using deep learning to classify biomedical images, and interpreting these deep-learned models is necessary for life-critical decisions and scientific discovery. Effective interpretation techniques accelerate biomarker discovery and provide new insights into the etiology, diagnosis, and treatment of disease. Most interpretation techniques aim to discover spatially-salient regions within images, but few techniques consider imagery with multiple channels of information. For instance, highly multiplexed tumor and tissue images have 30-100 channels and require interpretation methods that work across many channels to provide deep molecular insights. We propose a novel channel embedding method that extracts features from each channel. We then use these features to train a classifier for prediction. Using this channel embedding, we apply an interpretation method to rank the most discriminative channels. To validate our approach, we conduct an ablation study on a synthetic dataset. Moreover, we demonstrate that our method aligns with biological findings on highly multiplexed images of breast cancer cells while outperforming baseline pipelines.

## 1 Introduction

Highly multiplexed imaging provides data on the spatial distribution of dozens to hundreds of different protein and protein modifications in a tissue. This provides an unprecedented view into the cells and structures that comprise healthy and diseased tissue. As such, highly multiplexed imaging is emerging as a potentially breakthrough technology in translational research and clinical diagnosis. Examples of highly multiplexed imaging technologies include imaging mass cytometry (IMC) [5], multiplexed ion beam imaging (MIBI) [6], co-detection by indexing (CODEX) [3], and cyclic immunofluorescence (CyCIF) [8].

Each image can comprise 30 to 100 unique channels (that each correspond to the detection of a specific protein) with millions of cells, and so computational tools are essential for analysis. To interpret the outputs of computational tools and answer specific research and clinical questions, it is critical to know which image channels are informative. Even though there is research on interpretation techniques for natural images [14,15, 7,18], channel- or target-wise importance ranking interpretation techniques for highly multiplexed images do not yet exist.

We introduce a novel system to automatically identify informative channels in highly multiplexed tissue images and to provide interpretable and potentially actionable insight for research and clinical applications. The process is illustrated in Figure 1. What follows is a description of our system for the goal of identifying the most informative channels for assessing the tumor grade of highly multiplexed images [5]. We first encode each channel using the shared weights of a ResNet18 [4] backbone encoder. To obtain an interpretable representation, which we refer to as channel embedding, we use an embedding encoder. Then, we train a classifier to produce a probabilistic prediction for each tumor grade class. Finally, we measure each channel’s contribution to the tumor grade classification by applying an interpretation technique, Backprop [18], that backpropagates gradients to the channel embedding. In our experimental results, we demonstrate that our system outperforms conventional algorithms [20, 4,10] combined with interpretation techniques [15,18] on the informative channel identification task for assessing tumor grade. Moreover, the informative channels identified by our novel method align with findings from a single cell data analysis [5], even though our approach does not require single cell segmentation.

**Fig. 1:**
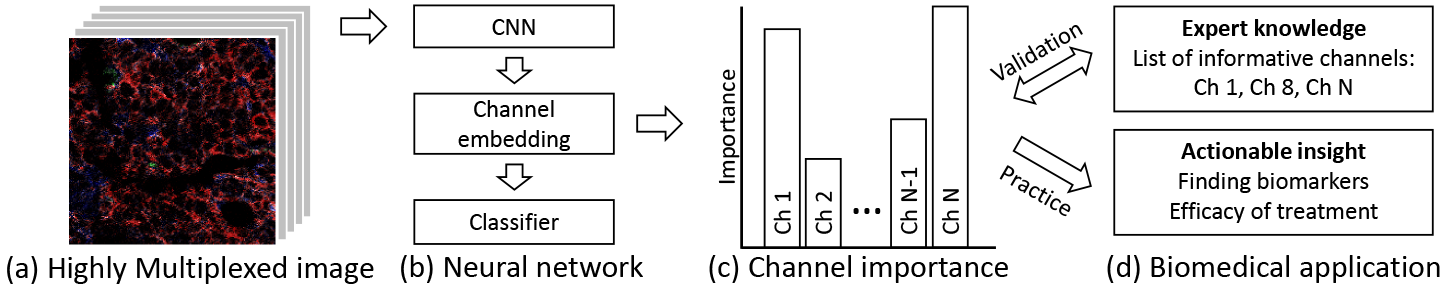
Informative channel identification. (a) Given highly multiplexed imaging data, we train (b) a neural network to encode a channel embedding and classify a label (*e.g*., tumor grade). Then, we measure (c) the classification task channel importance by adopting an interpretation method to the channel embedding. (d) We evaluate our system by comparing the predicted informative channels to expert knowledge, and provide new insights for clinicians and pathologists.

## 2 Related Works

### Interpretation Techniques for Neural Networks

One category of neural network interpretation is model-agnostic. Backprop [18] and Grad-CAM [15] backpropagate gradients to produce an attention map highlighting important regions in the image. Filter visualization techniques [12, 21,1] typically visualize the information extracted from filters. LIME [14], DeepLIFT [17], and SHAP [11] compute the contribution of each feature for a given example. TCAV [7] defines high-level concepts to quantify a model’s prediction sensitivity.

Another category is self-interpretable neural networks. SENN [13] trains a self-explaining model, which consists of classification and explanation branches. Zhang *et al*. [22] modify a traditional convolutional neural network (CNN) by adding masking layers followed by convolution layers to force activations to be localized. Building on this work, Zhang *et al*. [23] visualize a CNN’s decision making process by learning a decision tree for a pre-trained model. However, these methods are not directly applicable to highly multiplexed input images.

### Frame-level Action Localization in Videos

Discovering informative channels is similar to localizing frames with target actions in videos. Action localization finds frames of interest in an entire video. CDC [16] predicts per-frame confidence scores using 3D convolutional neural networks. BSN [10] and BMN [9] adopt 2D convolutions to estimate actionness, starting time, and ending time at each frame. These methods can be applicable to informative channel identification by using their per-channel classification as a measure of channel importance. However, since they perform prediction by classifying one channel at a time, their learned features may not be generalizable and consequently their resulting accuracy may not be sufficient.

## 3 Proposed Method

To enable the identification of informative channels, we propose a network architecture with a backbone encoder, an embedding encoder, and a classifier (Fig. 2). First, we will introduce the backbone and embedding encoders, which each extract features from each channel independently. For interpretation, we represent each input image channel as a single value *r_i_*. Next, we train a classifier that takes the channel embedding **r**, which is a concatenation of *r_i_* across channels. The classifier yields predictions, **p**. Once our network is trained, we apply Back-prop [18] to the channel embedding to produce an attention map, which is then used to rank channels in order of importance.

**Fig. 2:**
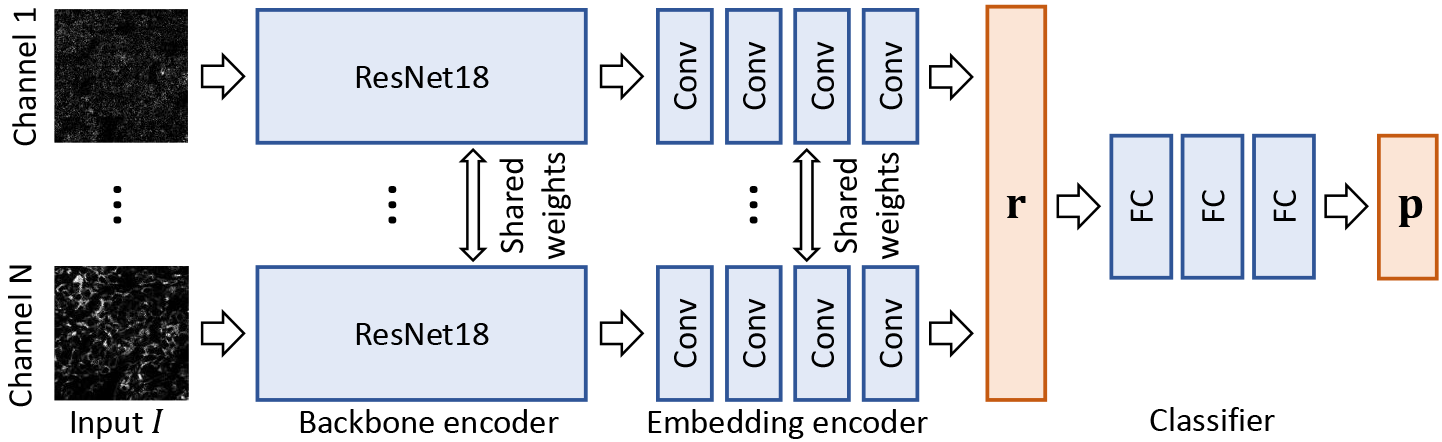
Our architecture. For an input highly multiplexed image *I*, we split channels and feed them into ResNet18 [4]. Next, the embedding encoder extracts an interpretable representation **r** = [*r*0,…,*r_N_*], where *N* is the number of channels. Both the backbone and embedding encoders share weights across channels. Finally, we adopt three fully-connected layers to estimate class probabilities **p**.

### 3.1 Channel Embedding

To leverage knowledge from previously-learned image classification tasks and to extract meaningful features from highly multiplexed images, we begin with ResNet18 [4] pretrained on ImageNet [2] as a backbone network. Naive application of ResNet18 does not allow us to identify channels of interest because it weights information across channels through its constituent convolutions (Fig. 3(a)). Even though 3D CNN [20] can suppress this problem by locally weighting channels, it still blends activations across adjacent channels (Fig 3(b)). Instead, we must extract features from each channel independently. However, doing so will require substantial memory resources (12M parameters per channel). To overcome these problems, we apply the same backbone encoder (shared weights) to each channel individually by modifying the first convolution layer of ResNet18 to accept a single-channel image as opposed to a three-channel (RGB) image (Fig 3(c)).

**Fig. 3:**
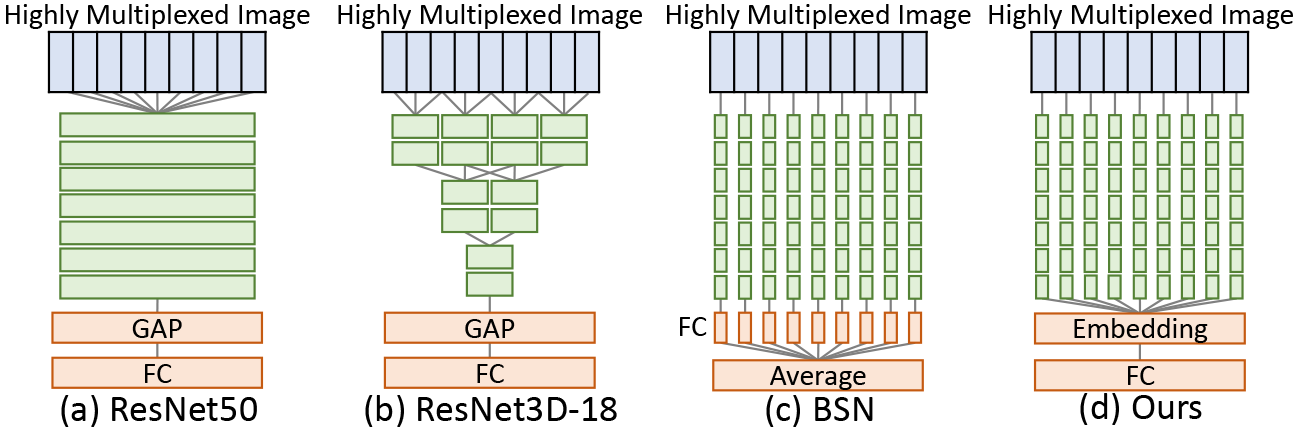
Architectures of baselines and our network. GAP and FC indicate a global average pooling and a fully connected layer. Unlike other baselines, we convert the input image into a channel embedding and then perform classification.

For interpretation, applying Backprop [18] to the backbone encoder output yields *N* 2D tensors, where *N* is the number of channels. The elements in each 2D tensor represent importance; however, these values are difficult to conceptually interpret. To avoid this issue, we add an embedding encoder that represents each channel as a single value *r_i_* (Fig 3(d)). The embedding encoder consists of three 3 × 3 and one 7 × 7 convolution layers with batch normalization and ReLU. The 3 × 3 and 7 × 7 convolution layers have 64 and 1 kernels, respectively. Concatenating the embedding encoder outputs yields a channel embedding **r** = [*r*_0_,…,*r_N_*].

To produce a prediction **p** for a classification task, we apply a classifier to the channel embedding **r**. We exploit two fully connected layers (FCs) with ReLU and one FC with softmax as a classifier. The first two FCs have 200 kernels.

### 3.2 Informative Channel Identification

After model training, we apply Backprop [18] to the channel embedding. Specifically, we first convert a given highly multiplexed image into a channel embedding and perform classification. We compute gradients at the channel embedding by backpropagating the gradients of the classification output. Then, we set the importance of each channel as the magnitude of its respective gradient. Unlike other systems using standard classification architectures [4, 20], our novel system yields a single value at each channel representing its importance in classification. Note that due to this design choice, our system can be used in a plug-and-play fashion with other interpretation methods [14,11,17] as well.

We measure the channel importance of all testing images and then average them across images to measure how informative each channel is for classification.

#### Implementation

We initialize weights in our network with random values except for the pre-trained ResNet18 backbone network. The spatial resolution of the input image *I* is 224 × 224 pixels. For data augmentation, we apply horizontal and vertical flips and random cropping. We use an Adam optimizer with a learning rate of 0.0001. The training process iterates for 100 epochs with early stopping while using a batch size of 32 and four Geforce Titan X GPUs.

## 4 Experimental Results

To validate the design of our model, we conduct an ablation study on a synthetic dataset classification task. To evaluate informative channel identification performance, we define a task in which modern deep neural networks [4, 20] achieve high accuracy. For a real-world application, we apply our model to a task of predicting tumor grade from a breast cancer dataset generated using IMC [5].

**Methods in Comparison:** To the best of our knowledge, there is no existing method for informative channel identification. As such, we implement baseline pipelines using modern classifiers, ResNet50 [4], ResNet3D-18 [20], and BSN [10], and model-agnostic interpretation methods, Backprop [15] and GradCAM [18]. We apply the interpretation techniques after training the classifiers to compare them with our system. While ResNet50 and ResNet3D-18 directly predict a class from the image, BSN classifies each channel separately and then aggregates the per-channel predictions to generate a single prediction. For ResNet50 and ResNet3D-18, we use Backprop [18] for interpretation. We convert each channel’s attention map into the channel importance by averaging it. For BSN, we adopt GradCAM [15] instead since BSN averages per-channel classification.

### 4.1 Synthetic Highly Multiplexed Image Classification

We build a synthetic dataset by emulating a highly multiplexed cell imaging environment. Generated images from this dataset contain a random number of dispersed circles of a fixed radius, whose intensities are sampled from bi-modal distributions (representing the signal arising when a target is either present or absent). We randomly choose two modes between [0.1,0.3] and [0.7, 0.9]. Each mode is randomly assigned a frequency of 0.2 or 0.8. We set the variance of intensities as 0.3. We add Gaussian noise to each image. For each ground truth, we randomly delegate two non-overlapping channels to associate with that ground truth. We synthesize 600 training, 300 validation, and 300 test images with 30 channels. Figure 4 visualizes three randomly-selected channels of a synthetic image compared to a real highly multiplexed image [5].

**Fig. 4:**
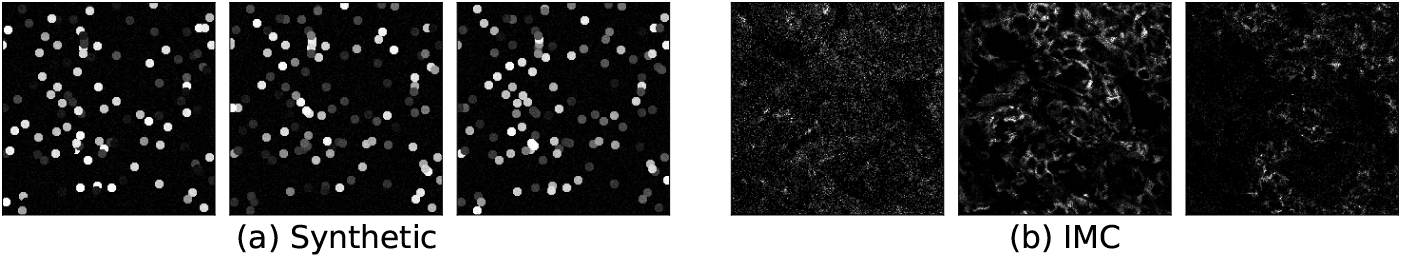
Sample channels from (a) synthetic and (b) IMC [5] images.

#### Evaluation Metric

To evaluate classification performance, we measure accuracy, which is the number of correct predictions divided by the number of total images. For the assessment of informative channel identification, we use Recall@*K*, which is a recall rate when a model proposes *K* most informative channels. Since there are six channels (two per class) associated with the classification, we set base *K* as 6 and expanded *K* as 10.

#### Ablation Study

We conduct an ablation study to find the best architecture design. Namely, we consider two choices for the backbone encoder: ResNet18 [4] and ResNet3D-18 [20]. For channel embedding, we consider two approaches: using shared weights across all channels or using independent channel-wise layers. Table 1 lists the scores of each setting. In terms of classification accuracy, all the settings are comparable. However, the purpose of this experiment is to examine the model’s ability to identify informative channels rather than classification. For informative channel identification, the ResNet18 + Shared setting achieves the best scores in terms of both Recall@6 and Recall@10. Since the backbone encoder, with 3D convolution layers, mixes information across channels, there is performance degradation in the ResNet3D-18 + Independent setting.

**Table 1:** Ablation study on the synthetic dataset.

| Settings |  | Scores |  |  |
| --- | --- | --- | --- | --- |
| Backbone | Embedding | Accuracy | Recall@6 | Recall@10 |
| ResNet18 | Independent | 98.0 | **83.3** | 83.3 |
| ResNet18 | Shared | 99.7 | **83.3** | **100.0** |
| ResNet3D-18 | Independent | **100.0** | 50.0 | 66.7 |
| ResNet3D-18 | Shared | 99.7 | 33.3 | 83.3 |

#### Results

Table 2 compares our system to the baselines. We find that our novel approach significantly outperforms the others in terms of Recall@6 and Recall@10 while achieving similar classification accuracy. This shows that our channel embedding is highly interpretable and effectively represents each channel.

**Table 2:** Quantitative comparison on the synthetic dataset.

| Representation | Method | #Param. | Acc. | Recall@6 | Recall@10 |
| --- | --- | --- | --- | --- | --- |
| Single prediction | ResNet50 [4] | 24M | **100.0** | 50.0 | 66.7 |
|  | ResNet3D-18 [20] | 33M | **100.0** | 50.0 | 50.0 |
| Channel-wise prediction | BSN [10] | 11M | 34.7 | 0.0 | 0.0 |
| Channel embedding | Ours | 12M | 99.7 | **83.3** | **100.0** |

### 4.2 Tumor Grade Classification

The breast cancer dataset [5] consists of multiplexed images collected using IMC. The images have 39 channels representing a set of proteins that are thought to be important for diagnosis or treatment. For each patient, clinical annotation is available. Here, we focus on identifying tumor grade: grade 1, grade 2, or grade 3. Tumor grade is an indicator of disease progression that is typically scored by a pathologist using only H&E images. We seek to identify the most informative channels among the 39 in the dataset with respect to the prediction of tumor grade. The network input is a multiplexed image and the output is a tumor grade. The dataset contains 723 tissue images from 352 breast cancer patients, and we split them into 506 training images and 217 test images. After training our network, we identify informative channels for predicting tumor grade.

#### Ground Truth Targets

Quantification of targets in single-cell analysis is correlated with clinical annotations such as tumor grade; however, this requires segmentation to isolate individual cells [5]. In contrast, we predict the tumor grade without using single-cell segmentation. Additionally, we demonstrate that our approach can interpret the importance of individual proteins (channels).

We use the single-cell averaged expression of the individual proteins to compare changes between grade 1, grade 2, and grade 3. Further, we use the sum of the absolute fold-change as a ground truth for analysis. Finally, to evaluate the pipelines, we use the same set of targets as those in the Jackson *et al*. study [5].

#### Evaluation Metric

For assessment of classification performance, we report the accuracy. To evaluate each model’s informative channel identification, we calculate the Spearman coefficients [19] using the ground truth. The Spearman coefficient measures the correlation between two lists of ranks.

#### Results

Table 3 compares our system to the baseline pipelines. Our classifier performs better than ResNet50 and BSN. For informative channel identification, our system significantly surpasses the baseline systems in Spearman coefficient. Figure 5 shows the importance of each channel predicted by our pipeline compared to the ground truth. We detect seven of the top 10 ground-truth targets, which are known proteins associated with tumor progression. One example is Ki-67, which represents a higher proliferation rate with increasing grade. Another example is CAIX, which represents a marker of hypoxia and also increases with a higher grade. Similarly, it was shown that low cytokeratins (like CK8/18, CK19 and anti-pan keratin (AE1)) are correlated with Grade 3 pathology [5].

**Table 3:**
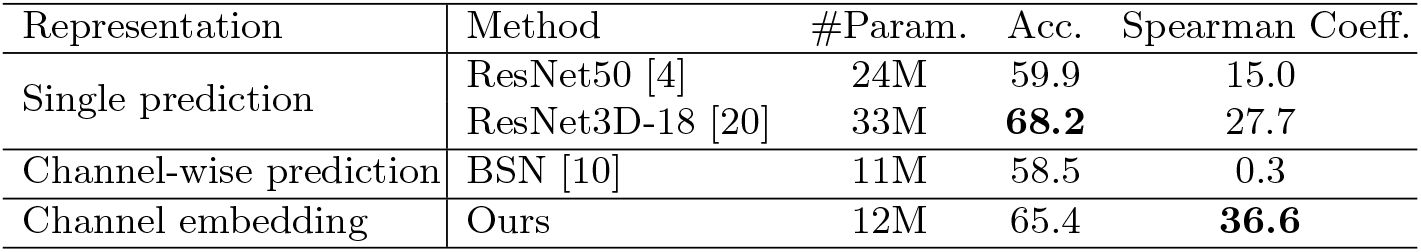
Quantitative results for tumor grading on the breast cancer dataset [5].

**Fig. 5:**
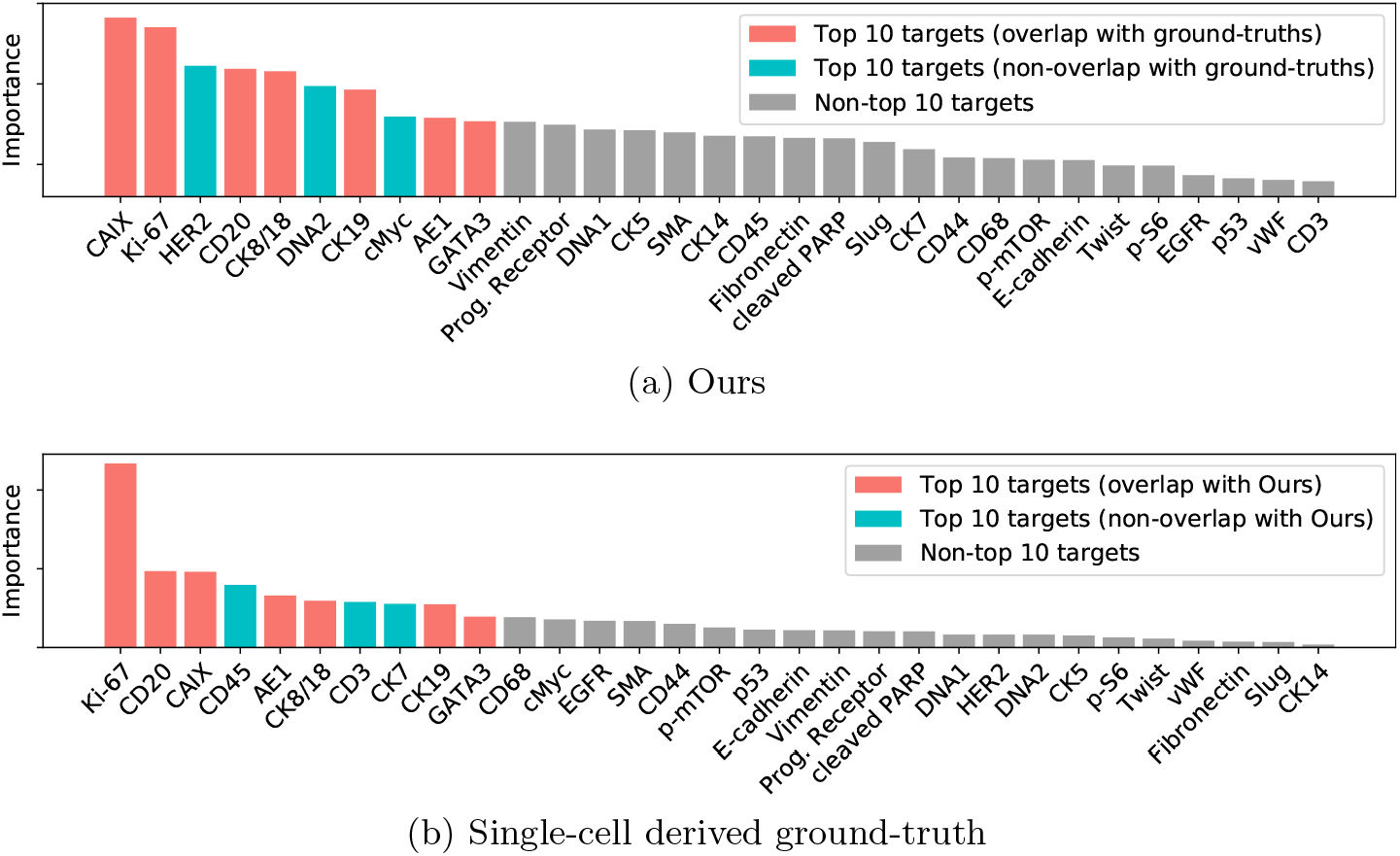
Measured target importance for tumor grade classification on the breast cancer dataset [5], ordered by importance. We highlight the top 10 targets.

## 5 Conclusions and Future Work

We have developed a novel pipeline for channel-wise importance interpretation. Our channel embedding effectively simplifies information in each channel while improving channel-wise interpretability. In the experimental results, we show that our pipeline outperforms existing methods [4,10, 20] in terms of classification and informative channel identification for tumor grade prediction [5].

Future work will focus on improving the approach and extending it to other datasets and prediction challenges including (i) biomarker discovery associated with survival time in breast cancer [5], (ii) discovery of cellular features predictive of treatment resistance in metastatic melanoma and other diseases, and (iii) the inclusion of spatial transcriptomic data.

## Acknowledgment

Hartland Jackson and Jana Fischer for help with the IMC dataset; This work was supported by NCI (U54-CA225088). Denis Schapiro was supported by an Early Postdoc Mobility fellowship (P2ZHP3_181475). Donglai Wei and Hanspeter Pfister have been partially supported by NSF grant NCS-FO (1835231). Salma Abdel Magid has been partially supported by Harvard University’s Graduate School of Arts and Sciences Graduate Prize Fellowship.

## Supplemental Materials

### 1 Calculation of Ground Truth Importance

We define a ground truth importance of a target *i* as

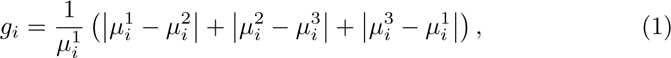

where 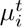 is the average intensity of target *i* across all single cells of a tumor grade *t*.

### 2 More Quantitative Results on the Synthetic Dataset

We report the Recall@*K* for different values of *K i.e*., different number of target proposals from an informative channel identification method. Table 1 compares informative channel identification methods in terms of Recall@*K*. It is observable that our method outperforms the other approaches for all values of *K*.

**Table 1:** Quantitative comparison on the synthetic dataset.

| Method | Recall@$K$ | | | | | | | |
| --- | --- | --- | --- | --- | --- | --- | --- | --- |
| | $K=6$ | $K=7$ | $K=8$ | $K=9$ | $K=10$ | $K=11$ | $K=12$ | $K=13$ |
| ResNet50 [4] | 50.0 | 50.0 | 66.7 | 66.7 | 66.7 | 66.7 | 66.7 | 83.3 |
| ResNet3D-18 [20] | 50.0 | 50.0 | 50.0 | 50.0 | 50.0 | 50.0 | 50.0 | 50.0 |
| BSN [10] | 0.0 | 0.0 | 0.0 | 0.0 | 0.0 | 0.0 | 0.0 | 0.0 |
| Ours | **83.3** | **83.3** | **100.0** | **100.0** | **100.0** | **100.0** | **100.0** | **100.0** |

### 3 Target Importance for Tumor Grade Classification

Fig. 1 shows the measured target importance for tumor grade classification on the breast cancer dataset [5]. While the conventional methods [4, 10, 20] combined with interpretation techniques [15, 18] detect only up to five targets among the top-10, seven targets identified by our pipeline overlap with the top-10 single cell derived ground-truth.

**Fig. 1:**
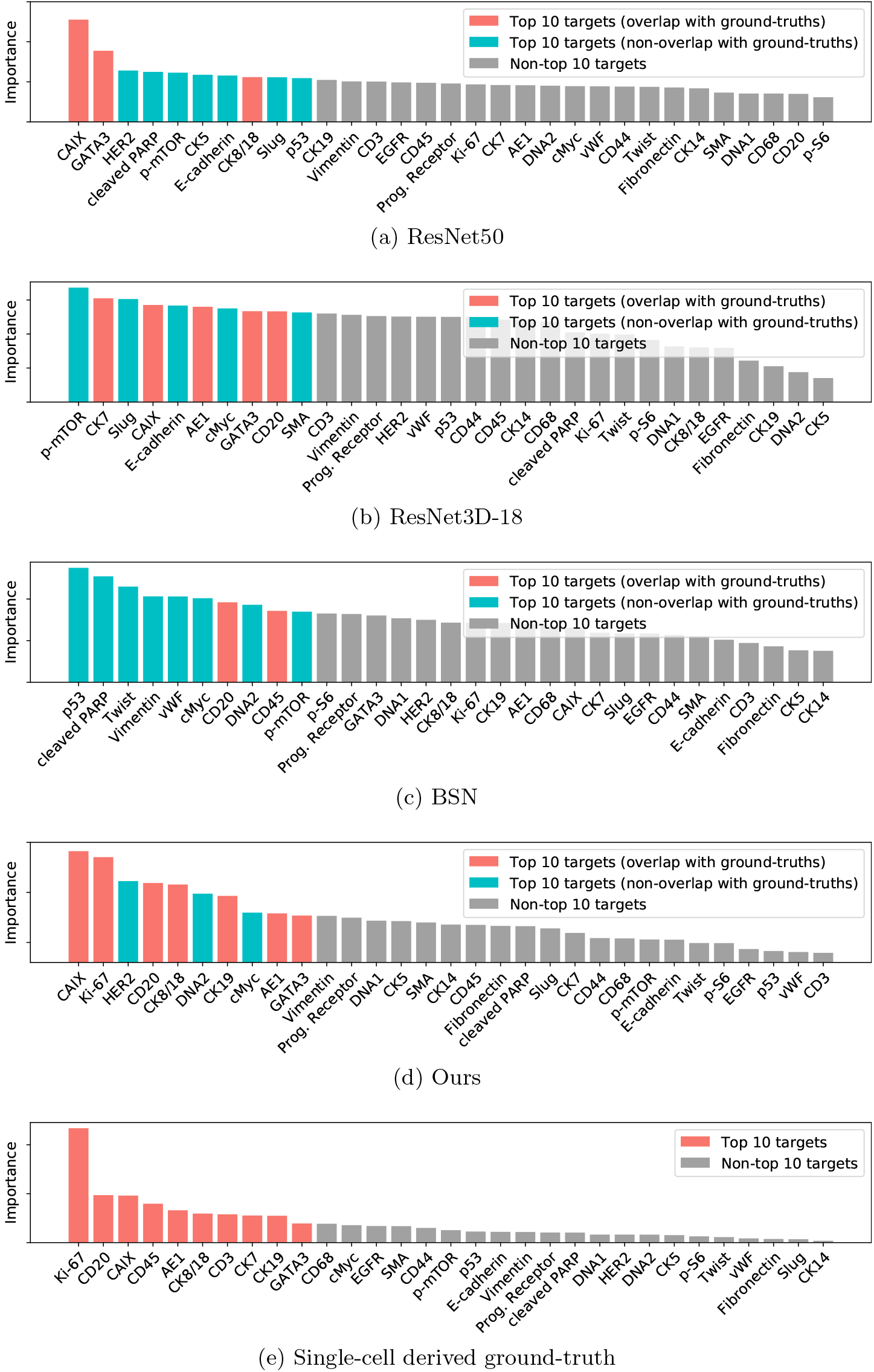
Measured target importance for tumor grade classification on the breast cancer dataset [5], ordered by importance. We highlight the top 10 targets.

